# 4G cloning: rapid gene assembly for expression of multisubunit protein complexes in diverse hosts

**DOI:** 10.1101/2024.06.17.599261

**Authors:** Michael Taschner, Joe Bradley Dickinson, Florian Roisné-Hamelin, Stephan Gruber

## Abstract

Multi-subunit protein complexes are at the heart of many cellular processes, and studying their biochemical activities and structures *in vitro* requires their reconstitution by recombinant expression and purification. Obtaining targets at sufficient purity and scale typically requires the screening of several protein variants and expression hosts. Existing cloning strategies allow to produce constructs for co-expression of proteins, but are often time-consuming, labour-intensive, host-specific, or involving error-prone assembly steps. Here we present a unique set of vectors together with a novel assembly strategy designed to overcome these limitations. It allows for the generation of expression constructs for multi-subunit protein complexes for various hosts in a single cloning step. Its modular nature allows the system to be easily extended to target additional expression hosts or to include new tags or regulatory sequences. As a proof of principle, we present the parallel construction of expression vectors for several Structural Maintenance of Chromosomes (SMC) complexes, allowing us to devise strategies for the recombinant production of these targets in bacteria, insect cells, and human cells, respectively. This work will help laboratories working on protein complexes to streamline their workflow, increase their productivity and improve the quality of the purified material.

## Introduction

Expression and purification of recombinant proteins is an integral part of many areas in molecular biology and biochemistry. Obtaining sufficient quantities of such samples at the desired level of purity frequently requires significant efforts in screening numerous expression constructs before finding one that gives satisfactory results.

Two decisions are crucial before starting to express and purify a new target. First, one has to select a suitable expression host (1). Bacterial expression in *E. coli* is usually the first choice due to the low cost, fast progress, minimal need for non-standard laboratory equipment, and ease of upscaling. Eukaryotic proteins, however, often require a more complex machinery for folding and/or post-translational modification and thus benefit from eukaryotic expression hosts. Second, an appropriate affinity tag needs to be found that allows for efficient purification of the target in a minimal number of steps without negatively impacting folding and function of the tagged protein.

For multi-subunit complexes the individual subunits can be expressed from separate vectors which are co-delivered into host cells, but this becomes inconvenient and unreliable for larger assemblies (>3 subunits). A better alternative is to produce multiple proteins from the same vector, with each subunit being expressed from its own gene expression cassette (GEC) with appropriate regulatory sequences. In recent years, several systems have been developed to facilitate the production of such multi-GEC plasmids for bacteria, insect cells, or mammalian cells (see for example (2–6). They are, however, host-specific and not easily cross-compatible. Moreover, cloning of constructs involves multi-step procedures that initially require the generation of single expression constructs (tagged or untagged) which are subsequently combined into larger assemblies. Modification to the expression strategies at a later time, such as changing tags or switching expression hosts, is often not straightforward and requires extensive new efforts. As an alternative to multi-GEC expression, several open reading frames (ORFs) can be expressed from a single cassette by connecting them via internal binding sites for the ribosome (‘RBS’ for bacteria; ‘IRES’ for eukaryotic cells) to create poly-cistronic constructs, or by a peptide linker of variable length to produce fusion proteins.

Traditional cloning methods (by restriction and ligation) are often labour-intensive, especially for multi-subunit expression vectors, as they typically require extensive handling and purification of DNA fragments (PCR purification, agarose gel-extraction, etc.), and re-sequencing of intermediates and final products if PCR techniques are employed. More recent developments offered convenient alternatives to these traditional approaches.

Golden Gate cloning uses Type IIS restriction enzymes, which cleave DNA outside their non-palindromic recognition sequence, to assemble one or more sequences provided in ‘donor’ vectors in the correct order into ‘acceptor’ vectors (7, 8). Donor and acceptor vectors differ in the placement and orientation of the Type IIS sites. Cloning is carried out in a ‘one-pot restriction/ligation’ reaction in which the desired circular plasmid (devoid of any recognition sites for the particular enzyme) accumulates. All fragments must have unique ‘sticky ends’ created by the Type IIS enzyme, and any internal restriction sites in target sequences have to be removed in a process termed ‘sequence domestication’. Due to the simple cut-and-paste mechanism without any PCR intermediate, the product does not need to be re-sequenced if the sequences of the acceptors and donors have already been adequately verified.

Gibson assembly (9) allows the construction of a target sequence from several linear fragments into a linearized vector in a short isothermal reaction, using short (15-25 bp) stretches of homology at the desired junctions. Fragments are often generated by PCR, and the product thus requires sequence-verification. Even if the fragments are created by error-free means, the assembly process involves the action of a DNA polymerase 100-200 bp around fragment junctions (9), so at least these areas need to be sequence-verified if they contain critical elements.

Both Golden Gate assembly (e.g. (4, 6)), Gibson assembly (e.g.(3)), or a combination thereof (10) have been successfully employed to create multi-GEC plasmids, but they either (i) use a step-wise cloning approach over multiple days which often requiring tedious DNA fragment purification and handling steps, (ii) involve intermediate PCR steps and thus require the large final product to be sequence verified, and/or (iii) are specifically designed to target only one particular expression host.

Here we present a new cloning strategy, which we term ‘**G**olden **G**ate-**g**uided **G**ibson Assembly’ or ‘4G cloning’ for short to produce complex expression vectors for several expression hosts. It solves the aforementioned problems and allows quick and reliable assembly of many construct variations in parallel. We provide a set of vectors that can easily be expanded to include new tags or regulatory sequences. For expression in insects and mammalian cells, the vectors are fully compatible with the recently described biGBac method (3). Our system allowed us to create several bacterial expression constructs for screening of the hexameric Smc5/6 complex from *Saccharomyces cerevisiae*, as well as similar constructs for the *Schizosaccharomyces pombe* and *Homo sapiens* Smc5/6 hexamers for expression in insect and mammalian cells, respectively. Moreover, the strategy has been successfully applied to produce the three-subunit Wadjet SMC complex for biochemical reconstitution and structural analysis. Lastly, although our work presented here is focused on the production of vectors for protein expression, the same assembly pipeline (4G-cloning) can be applied to other areas requiring quick modular assembly of repeating building blocks.

## Results

### Assembly of multi-GEC expression plasmids using *G*olden *G*ate-*g*uided *G*ibson Assembly

Here, we devised a cloning strategy based on Gibson assembly of Golden Gate-customized gene expression cassettes (GECs), producing final multi-gene expression vectors from sequence-verified elements in a single cloning step (**Fig. 1**) (‘**G**olden **G**ate-**g**uided **G**ibson assembly’ or 4G cloning). Briefly, Golden Gate assembly of compatible DNA fragments first produces a linear product, the GEC. Multiple GECs with appropriate terminal overhangs are then directly mixed and inserted into an acceptor vector in a defined order by Gibson assembly. We prepared a DNA-element library (Table 1), with which production of multi-GEC expression plasmids can be carried out in a single day with minimal hands-on time. First, Golden Gate reactions are set up individually for each GEC including donors for all desired elements (**Fig. 1**, top half), with complete flexibility regarding inclusion of a tag, host-specific regulatory sequences, and position of the GEC in the final construct. Without further amplification or purification, the products of these reactions are then combined and mixed with linearized acceptor vector to produce the desired plasmid by Gibson assembly on the same day (**Fig. 1**, bottom half). Upon transformation into *E. coli*, positive clones can be identified and then used directly for expression screening without further sequencing, because GECs are assembled from sequence-verified parts by Golden Gate assembly, and any mutation introduced during Gibson assembly is limited to neutral spacer sequences surrounding the junction sites.

**Table 1:** Plasmids for 4G cloning

|  | plasmid name | resistance | strain | description |
| --- | --- | --- | --- | --- |
| Acceptors | pMulti_A_ccdB | Amp | ccdB surv. | GEC acceptor for *E. coli* expression, Ampicillin-resistance |
|  | pMulti_K_ccdB | Kan | ccdB surv. | GEC acceptor for *E. coli* expression, Kanamycin-resistance |
|  | pMulti_S_ccdB | Strep | ccdB surv. | GEC acceptor for *E. coli* expression, Streptomycin-resistance |
| Promoter/Terminator donors | pD(P+T)_T7 | Cm | pir | Promoter/terminator for *E. coli* expression, no tag |
|  | pD(P+T)_T7N | Cm | pir | Promoter/terminator for *E. coli* expression, N-term. tag |
|  | pD(P+T)_T7C | Cm | pir | Promoter/terminator for *E. coli* expression, C-term. tag |
|  | pD(P+T)_ph | Cm | pir | Promoter/terminator for insect expression, no tag |
|  | pD(P+T)_phN | Cm | pir | Promoter/terminator for insect expression, N-term. tag |
|  | pD(P+T)_phC | Cm | pir | Promoter/terminator for insect expression, C-term. tag |
|  | pD(P+T)_CMV | Cm | pir | Promoter/terminator for mammalian expression, no tag |
|  | pD(P+T)_CMVN | Cm | pir | Promoter/terminator for mammalian expression, N-term. tag |
|  | pD(P+T)_CMVC | Cm | pir | Promoter/terminator for mammalian expression, C-term. tag |
| Gibson overhang donors | pD(Gib)_5'alpha | Cm | pir | Gibson overhang , adds alpha-sequence to 5'-end |
|  | pD(Gib)_5'beta | Cm | pir | Gibson overhang , adds beta-sequence to 5'-end |
|  | pD(Gib)_5'gamma | Cm | pir | Gibson overhang , adds gamma-sequence to 5'-end |
|  | pD(Gib)_5'delta | Cm | pir | Gibson overhang , adds delta-sequence to 5'-end |
|  | pD(Gib)_3'beta | Cm | pir | Gibson overhang , adds beta-sequence to 3'-end |
|  | pD(Gib)_3'gamma | Cm | pir | Gibson overhang , adds gamma-sequence to 3'-end |
|  | pD(Gib)_3'delta | Cm | pir | Gibson overhang , adds delta-sequence to 3'-end |
|  | pD(Gib)_3'omega | Cm | pir | Gibson overhang , adds omega-sequence to 3'-end |
| N-terminal tag donors | pD(Nt)_His6 | Cm | pir | N-terminal hexa-histidine, non-cleavable |
|  | pD(Nt)_His6-TEV | Cm | pir | N-terminal hexa-histidine, TEV-cleavable |
|  | pD(Nt)_His6-3C | Cm | pir | N-terminal hexa-histidine, 3C-cleavable |
|  | pD(Nt)_His6-FLAG3 | Cm | pir | N-terminal hexa-histidine + triple-FLAG, non-cleavable |
|  | pD(Nt)_His6-FLAG3-TEV | Cm | pir | N-terminal hexa-histidine + triple-FLAG, TEV-cleavable |
|  | pD(Nt)_His6-HA2-TEV | Cm | pir | N-terminal hexa-histidine + double-HA, TEV-cleavable |
|  | pD(Nt)_His6-MBP-TEV | Cm | pir | N-terminal hexa-histidine + MBP, TEV-cleavable |
|  | pD(Nt)_His6-GST-TEV | Cm | pir | N-terminal hexa-histidine + GST, TEV-cleavable |
|  | pD(Nt)_His6-SUMO | Cm | pir | N-terminal hexa-histidine + SUMO, Senp2-cleavable |
|  | pD(Nt)_GFP | Cm | pir | N-terminal GFP, non-cleavable |
|  | pD(Nt)_GFP-TEV | Cm | pir | N-terminal GFP, TEV-cleavable |
|  | pD(Nt)_GFP-3C | Cm | pir | N-terminal GFP, 3C-cleavable |
|  | pD(Nt)_TS-3C | Cm | pir | N-terminal Twin-Strep, 3C-cleavable |
|  | pD(Nt)_His10-TS-3C | Cm | pir | N-terminal deca-histidine + Twin-Strep, 3C-cleavable |
|  | pD(Nt)_HALO | Cm | pir | N-terminal HALO, non-cleavable |
| C-terminal tag donors | pD(Ct)_His8 | Cm | pir | C-terminal octa-histidine, non-cleavable |
|  | pD(Ct)_3C-His8 | Cm | pir | C-terminal octa-histidine, 3C-cleavable |
|  | pD(Ct)_3C-GFP | Cm | pir | C-terminal GFP, 3C-cleavable |
|  | pD(Ct)_3C-Venus-His8 | Cm | pir | C-terminal octa-histidine + Venus, 3C-cleavable |
|  | pD(Ct)_CPD-His10 | Cm | pir | C-terminal deca-histidine + CPD, phytate-cleavable |
|  | pD(Ct)_TS | Cm | pir | C-terminal Twin-Strep, non-cleavable |
|  | pD(Ct)_3C-TS | Cm | pir | C-terminal Twin-Strep, 3C-cleavable |
|  | pD(Ct)_3C-TS-His10 | Cm | pir | C-terminal deca-histidine + Twin-Strep, 3C-cleavable |
|  | pD(Ct)_Avi-3C-TS | Cm | pir | C-terminal AviTag + Twin-Strep, partially 3C-cleavable |
|  | pD(Ct)_FLAG | Cm | pir | C-terminal FLAG, non-cleavable |
|  | pD(Ct)_FLAG3 | Cm | pir | C-terminal triple-FLAG, non-cleavable |
|  | pD(Ct)_MYC9 | Cm | pir | C-terminal nona-myc, non-cleavable |
|  | pD(Ct)_PK6 | Cm | pir | C-terminal hexa-V5, non-cleavable |
|  | pD(Ct)_HALO | Cm | pir | C-terminal HALO, non-cleavable |

Amp: Ampicillin, Kan: Kanamycin, Strep: Streptomycin, MBP: Maltose-Binding-Protein, GST: Glutathione-S-Transferase, GFP: Green Fluorescent Protein, CPD: Cysteine-Protease-Domain

**Figure 1:**
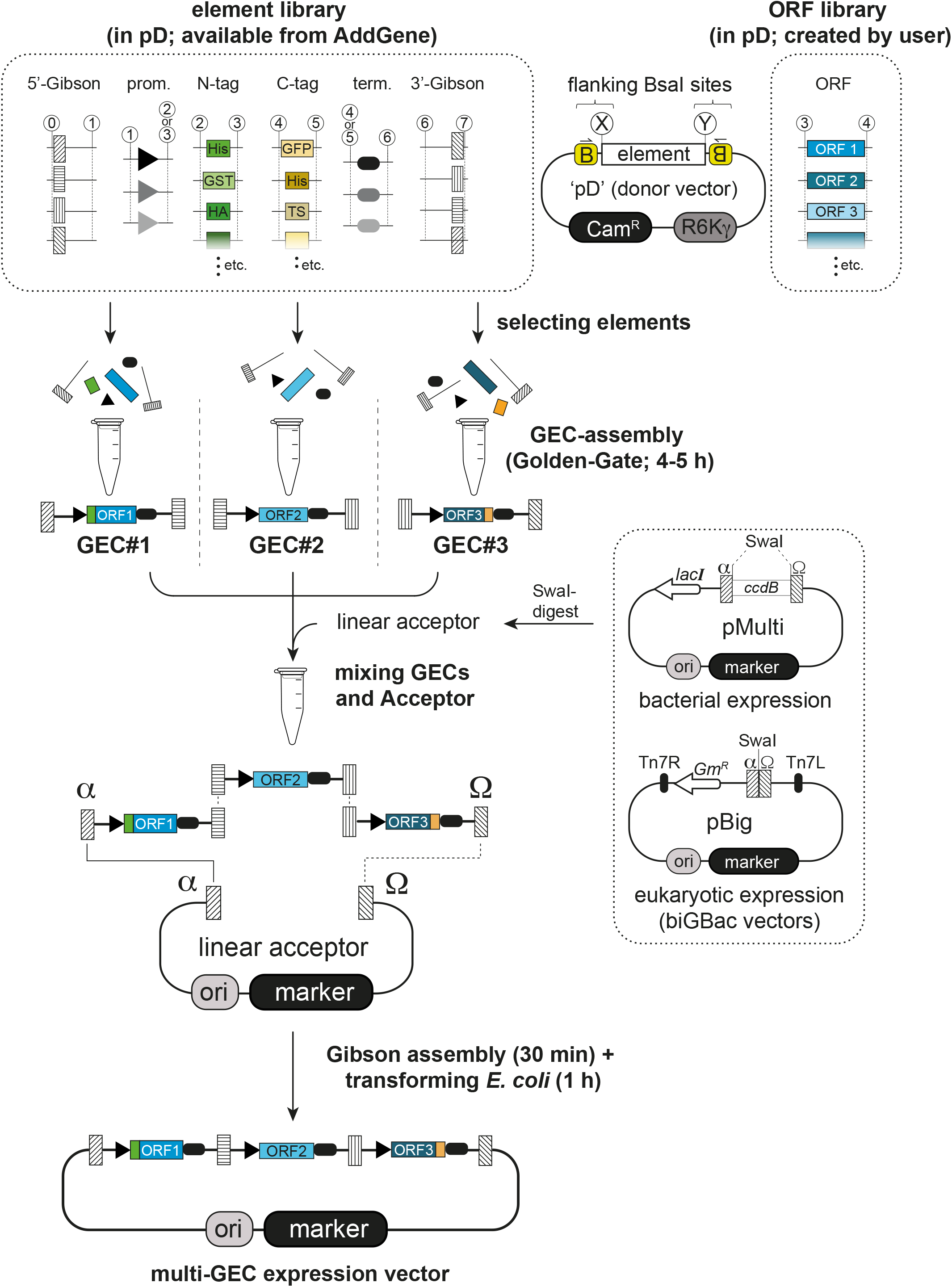
Overview of the ‘Golden Gate-guided Gibson (4G cloning) procedure. In a first step, individual linear gene expression cassettes (GECs) are assembled from individual smaller elements by Golden Gate cloning. These elements are excised from individual circular donor vectors (‘pD’; see generic representation next to the element library box), giving specific sticky ends depending on the element type (numbers in white circles). Multiple GECs with compatible homology ends from such reactions are then directly inserted into dedicated linearized acceptor vectors by Gibson assembly without further isolation/purification. As a result, a multi-GEC expression plasmid can be created in a single day from prepared donors.

We started by creating the building blocks, here designated as “elements”, for GEC assembly in a set of Golden Gate donor plasmids. Examples are shown schematically in **Fig. S1A**, and a more detailed view is provided in **Fig. S1D-F**. Each donor plasmid contains one or two elements (such as an ORF-element, a TAG-element, promoter-elements and terminator-elements, or Gibson-elements) (**Fig. S1 and Table 1**), which include appropriate flanks for subsequent Golden Gate assembly (see Materials and Methods). The flanks harbor recognition sites for the enzyme BsaI, cleavage of which creates unique 4 bp ‘sticky ends’ (numbers/letters in white circles in all figures). The cloning of these elements into donor plasmids by PCR and Gibson assembly (**Fig. S1B**) also allows for sequence verification and for concomitant sequence domestication, i.e. the removal of any internal BsaI sites in target sequences (**Fig. S1C**). We have successfully removed up to five internal BsaI sites in a single step in this way. All donors have a backbone of 1833 bp containing a chloramphenicol resistance marker (Cam^R^) and a conditional origin of replication (R6Kγ) only functional in bacteria expressing the *pir* gene (**Fig. 1, top right**).

Only donors for ORF-elements need to be created and sequence-verified by the user for each new expression target, i.e. each subunit of a protein complex of interest. The remaining set of vectors (**Table 1**) will be made available from AddGene. Donors for N- and C-terminal TAG-elements carry inserts that can be fused with the 5’ or the 3’ end of an ORF-element, respectively. We created such TAG-elements for many commonly used protein purification tags, with several of them including protease recognition sites for optional tag removal (TEV or 3C; **Table 1**), but new donors for any additional tag-elements can be quickly created using the same strategy (**Fig. S1B**,**C**). Upstream and downstream sequences are provided by promoter-elements and terminator-elements, generally combined in ‘P+T’-donors. We created such P+T combinations for expression in bacteria (T7 promoter; T7 terminator), in insect cells (polyhedrin promoter; SV40-polyA), and in mammalian cells (CMV promoter; bGH-polyA). Importantly, the P+T donors each exist in 3 forms to allow untagged expression or addition of a N-or C-terminal tag-element (**Fig. S1A, Fig. S1E**, and **Table 1**). Lastly, donors for 5’-and 3’-Gibson-elements provide sequences that attach upstream of the promoter element and downstream of the terminator element, respectively (**Fig. S1A** and **Fig. S1F**). They add terminal homology regions of around 20 bp for Gibson assembly which are separated from the internal elements of the GEC by 200-300 bp long spacers (to avoid sequence alterations in critical parts during Gibson assembly).

We also built acceptor vectors for the final multi-GEC assemblies allowing for expression in bacteria. Three pMulti-plasmids differing in their antibiotic resistance marker (Ampicillin, Kanamycin, or Streptomycin; see box on the right in **Fig. 1**) contain a *ccdB* suicide cassette to avoid vector-background during cloning, and thus are maintained in appropriate *ccdB*-resistant strains. Digestion with the enzyme SwaI removes the suicide cassette and exposes homology regions (α and Ω) used for GEC insertion by Gibson assembly. For creating multi-GEC plasmid for insect and mammalian expression (see below) we use plasmids pBig1a-pBig1e from the biGBac system (available from AddGene, plasmid kit #1000000088) which are linearized in the same way. These vectors are compatible because our α and Ω overhangs are identical to those described originally for the biGBac system (3).

Correct formation of the desired multi-GEC vectors depends on the complete assembly and subsequent joining of linear GECs. We reasoned that the efficiency would decrease with increasing number of GECs and set out to test this. We determined the efficiencies of obtaining clones containing either one, two, three, or four GECs by counting the number of colonies after transformation of the 4G cloning reaction (see **Fig. S1G** for an overview of the assembly process). As expected, the total number of clones decreased progressively from >4000 for a single GEC to around 150 for four GECs (**Fig. 2**, top). We then randomly picked five colonies for each plate, isolated the candidate plasmid, and analyzed it by restriction digest. While the accuracy of assembly decreased with increasing number of GECs, 40 % (2/5) of the picked clones with four GECs were still correct (**Fig. 2**, bottom). Of note, to allow simultaneous assembly of even more GECs into a single vector, additional Gibson-element donors will have to be designed.

**Figure 2:**
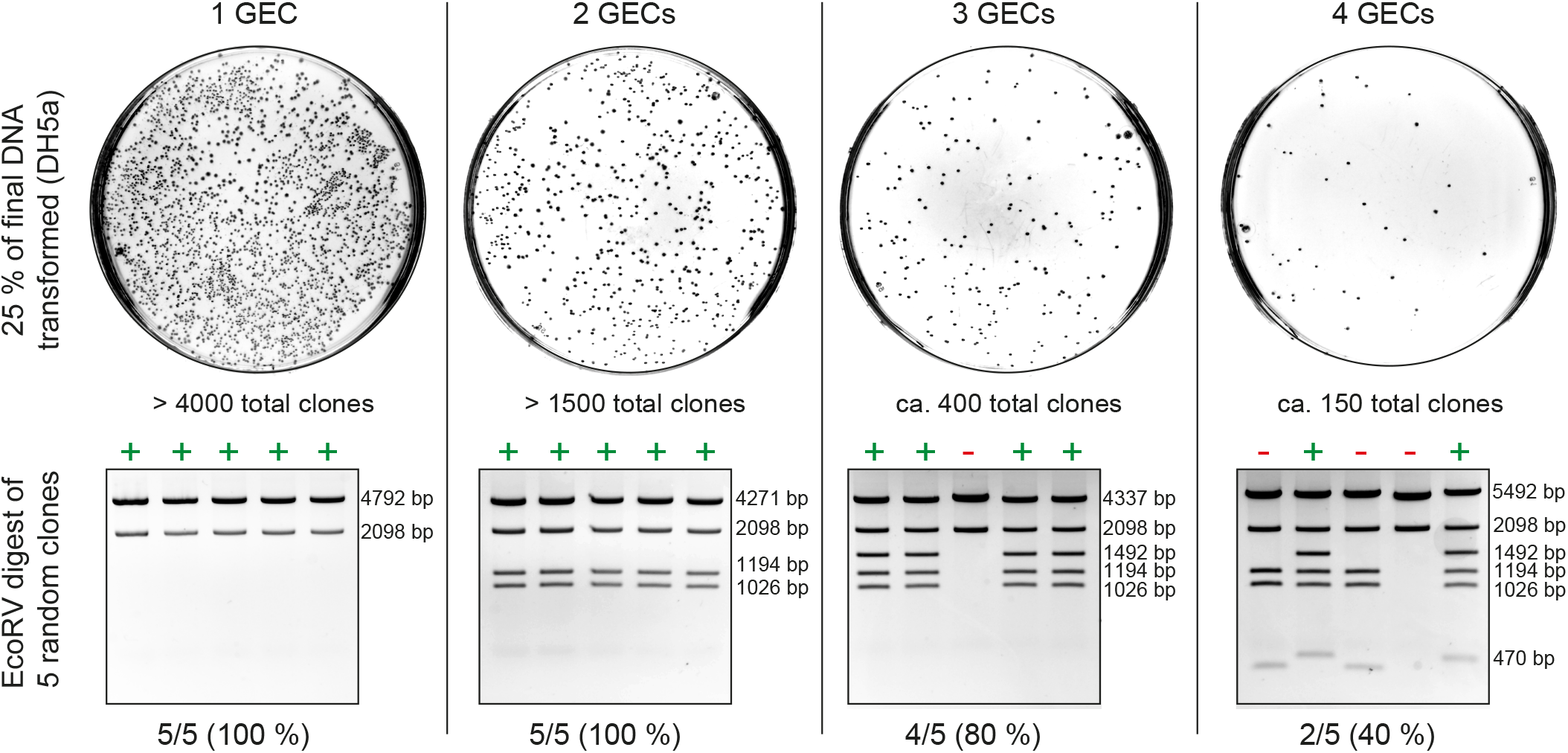
Test of assembly efficiencies with up to four GECs. 25 % of the final Gibson assembly reaction (2.5 µl out of 10 µl) were transformed into competent cells and the total number of colonies was estimated. Plasmids were isolated from 5 randomly chosen clones and tested by restriction digest. The numbers on the right side of the gel indicates the expected fragment sizes of positive clones.

Due to the quick and easy assembly process, 4G cloning allows to create many versions of expression plasmids in parallel. For example, if the best position for a purification tag is unknown, three GEC versions for each subunit (untagged, with N-terminal tag, or with C-terminal tag) can be prepared and then combined in a way to screen all possibilities without significantly increasing the workload (see below). Also, if a certain expression host (like *E. coli*) turns out to be a poor choice for production of a given target, similar vectors for a different host can be created very quickly and with minimal adjustments (i.e. usage of different promoter- and terminator-elements and acceptor vectors). We have recently reported the successful application of this cloning strategy for the expression of two distinct multi-subunit protein complexes in *E. coli* (11–14). Here we briefly describe the initial creation and further development of these expression tools based on a challenging target complex.

### Expression of the hexameric Smc5/6 complex from budding yeast in *E. coli*

Structural maintenance of chromosomes (SMC) complexes regulate essential aspects of chromosome structure and segregation in all domains of life, and share a common architecture containing two SMC proteins and several non-SMC components (15, 16). Eukaryotes contain three distinct SMC complexes called cohesin, condensin, and Smc5/6, with prominent roles in sister chromatid cohesion, chromosome condensation, and DNA repair, respectively. Whereas recombinant production of active SMC complexes has been achieved using eukaryotic expression hosts, the obtained yields and purity is limited, particularly for the hexameric yeast Smc5/6 complexes. At its core this complex contains the SMC proteins Smc5 and Smc6, as well as four non-SMC-elements (Nse1-Nse4) (17). Evidence in the literature indicates that several of these proteins from various species can be produced as smaller sub-complexes in *E. coli* (18, 19).

We set out to test production of the hexameric holo-complex from budding yeast in *E. coli* using 4G cloning; we prepared the six necessary ORF-elements by PCR amplification of the respective coding sequences from the yeast genome, followed by Gibson assembly into the linear donor vector according to **Fig. S1B** and **C**. Internal BsaI sites for Nse2, Smc5, and Smc6 were mutated as shown in **Fig. S1C**. Having no prior knowledge about the optimal type and position of affinity tag for this complex, we decided to test all possible 12 positions for a single TwinStrep tag-element in an unbiased way. An overview of our cloning procedure is shown in **Fig. 3A** and **Fig. S2**. We first assembled three GECs with T7 promoter-element and T7 terminator-element for each subunit (for untagged as well as N-or C-terminally Twin-Strep-tagged versions), giving a total of 18 GECs. Upstream and downstream homology-elements were chosen to create vectors with three GECs (Nse4/Nse3/Nse1, and Smc6/Smc5/Nse2). Golden Gate products were mixed in several combinations with linearized pMulti plasmids to give 14 final expression vectors. The Nse4/Nse3/Nse1 combinations were inserted into pMulti_K (with Kan^R^), while the Smc6/Smc5/Nse2 combinations were inserted into pMulti_A (with Amp^R^). Seven plasmids for each GEC-combination were created, one with all subunits untagged, and six for all possible tag positions (see **Fig. S2** for a detailed view). Importantly, all these vectors were created in parallel in a single day. 13 combinations of these plasmids (12 for the possible tag positions and one with all untagged subunits as negative control for purification) were then co-transformed into *E. coli* for small scale test expressions and Strep-Tactin pulldowns. The result of this experiment is shown in **Fig. 3B**. Minimal background was observed after Strep-Tactin pulldowns using extracts of cells expressing the untagged complex (lane 1). Tagging of Nse1, Nse2, Nse3, Nse4, or Smc5 gave similar results, regardless of whether the TwinStrep-tag was added to the N- or the C-terminal end of the respective protein. These pulldowns yielded high amounts of material, but the band for Smc6 was always clearly weaker than the bands for the other subunits (lanes 2-11), indicating poor subunit stoichiometry. When the tag was placed at the N-terminus of Smc6 the amount of protein obtained in the pulldowns was minimal (lane 12), indicating that Smc6 does not tolerate this tag position. Moving the tag to the Smc6 C-terminus, however, yielded material after pulldown. While the overall yield here (lane 13) was lower than with tags on any of the other subunits (lanes 2-11), the resulting complex appeared to have a balanced subunit stoichiometry.

**Figure 3:**
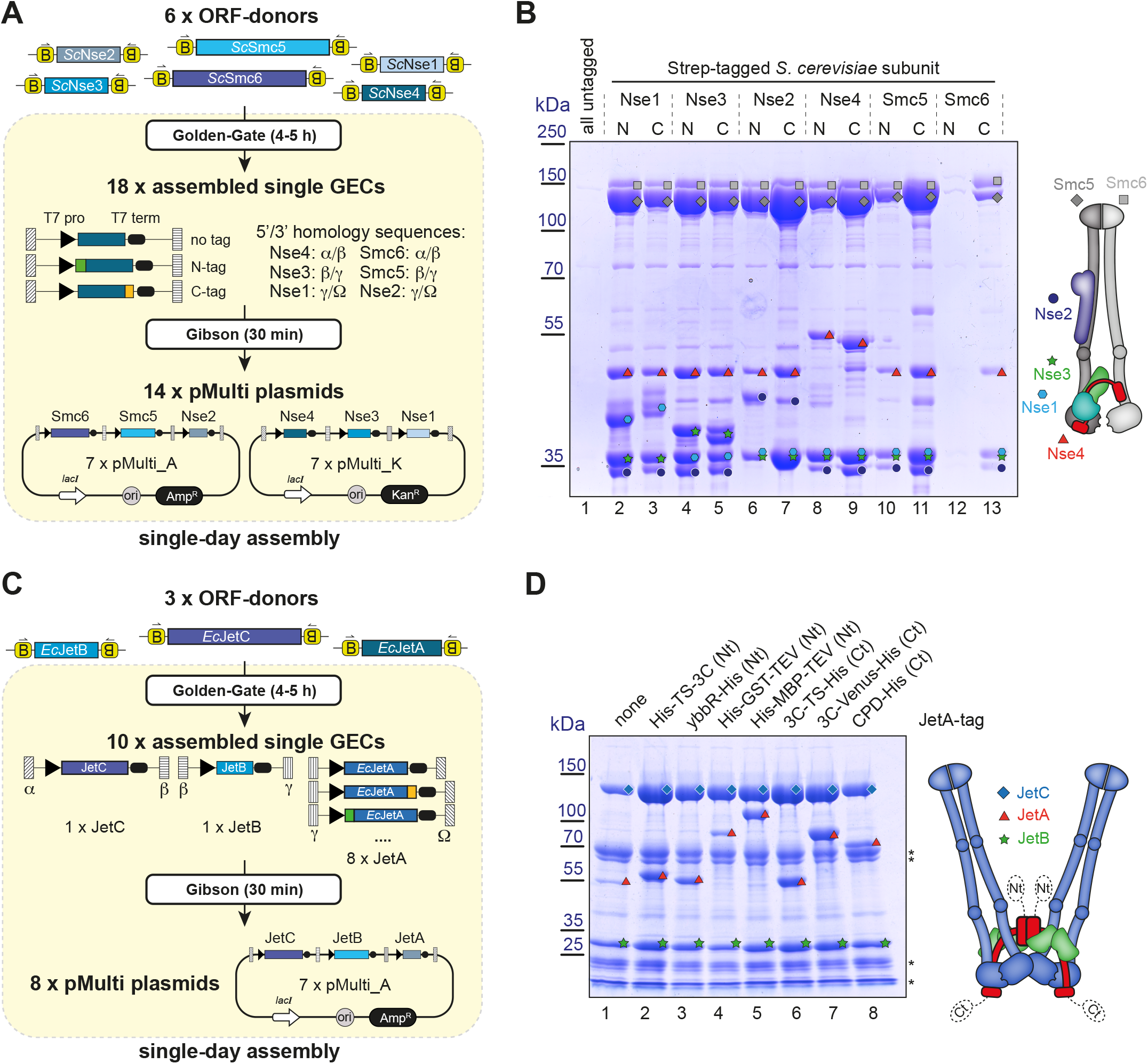
(A) Overview of the cloning procedure to produce vectors for expression of the *S. cerevisiae* Smc5/6 complex in *E. coli*. The six individual ORF donors for Smc6, Smc5, Nse4, Nse3, Nse2, and Nse1 were used to create 18 GECs (each subunit either untagged, or N- or C-terminally tagged) containing T7 promoters and terminators as well as dedicated Gibson overhangs to specify their position in the final vector. Subsequent Gibson-Assembly into pMulti_A or pMulti_K resulted in 14 vectors which were used to obtain 13 variations regarding the placement of the affinity tag (see also Fig. S2).(B) Coomassie-stained SDS-gel showing the results of the expression screen for the *S. cerevisiae* Smc5/6 complex in *E. coli*. Only a C-terminal tag on Smc6 gives a stoichiometric complex after a single affinity step. The scheme on the right shows the overall architecture of the hexameric complex. Colored symbols for each subunit are used to identify the bands in the gel. (C) Overview of the cloning procedure to produce vectors for expression of the *E. coli* JetABC complex. The three individual ORF donors were used to create eight pMulti_A vectors differing in the tag added to the JetA subunit on the third position.(D) Coomassie-stained SDS-gel showing the results of the expression screen for *E. coli* JetABC. Differences in yield can be observed when comparing different tags added to the JetA subunit. The untagged complex also has a certain affinity for the purification resin (lane 1). Additional bands represent contaminants and/or degradation products of subunits.

### Tag-screening for the JetABC complex from *E. coli*

Bacteria have evolved sophisticated defence systems to fight incoming phages and so-called ‘selfish’ DNA elements. One of them, ‘Wadjet’/JetABCD (also called MksBEFG/EptABCD) restricts plasmid transmission. It encodes for a sensor component, JetABC, which is an SMC complex resembling the bacterial condensin MukBEF, associated with the effector JetD, a TOPRIM domain-containing nuclease (12, 20, 21). JetABC forms a dimer of motor units that identifies smaller circular DNAs by loop-extrusion that are then cleaved by the JetD nuclease (12, 14, 20, 22). We previously expressed the JetABC complex with an N-terminal ‘His-TwinStrep-3C’-tag on the N-terminus of the kleisin subunit JetA (12), after screening for the best tag position using a strategy similar to the one described for Smc5/6. To screen other tags on the same subunit for specific applications (i.e. complex labelling) or for increased yield, we generated eight expression constructs in parallel by 4G cloning (**Fig. 3C**). GECs for JetC, JetB, and JetA were created, with JetA being left either untagged or receiving four N- or three C-terminal tag combinations that all contain a His-tag for direct comparison by Ni^2+^-NTA pulldown. The cassettes were inserted into the pMulti_A vector and transformed into *E. coli* for expression tests. Bands for all 3 subunits (together with additional bands representing contaminants and/or degradation products) were observed even after pulldown of the untagged complex (**Fig. 3D**, lane 1), consistent with a previous report that the JetC subunits binds to the His-affinity resin in a non-specific manner (22). The yield here was, however, clearly lower than for the other constructs containing a tag. Among those some clearly produced more protein than others, with the previously described construct (**Fig. 3D**, lane 2) being among the options with superior yield.

### 4G cloning for expression in eukaryotic hosts

The above results demonstrate that quick cloning of multi-subunit assemblies for expression in *E. coli* is feasible with 4G cloning, even for challenging targets like eukaryotic SMC complexes. However, we failed to express and purify Smc5/6 complexes from *S. pombe* and *H. sapiens* in *E. coli*, apparently due to limited expression of the SMC subunits. To solve this issue, we shifted our attention to expression in eukaryotic hosts (insect and mammalian cells). We created promoter- and terminator-elements for expression in these hosts (polyhedrin promoter/SV40 polyA or CMV promoter/bGH polyA; respectively) and performed 4G cloning with the respective ORF-elements cloned for *S. pombe* and *H. sapiens* Smc5/6 subunits. The combinations Smc6/Smc5/Nse2 and Nse4/Nse3/Nse1 were cloned into pBig1a and pBig1b, respectively, then combined into pBig2 according to the published procedure (3), and a similar tag-screening procedure as described for the budding yeast complex was carried out (**Fig. S3**). The *S. pombe* subunits were put under the control of insect cell promoters and the final vectors were used to produce baculoviruses which were tested for protein expression in small scale. The results shown in **Fig. 4A** show that stoichiometric complexes can be obtained in a single purification step when tagging the SMC subunits, regardless of whether the tag is on the N- or the C-terminus. Tagging any of the smaller Nse subunits consistently yields excess of the small subunits, either of the Nse4/3/1 sub-complex or of Nse2.

**Figure 4:**
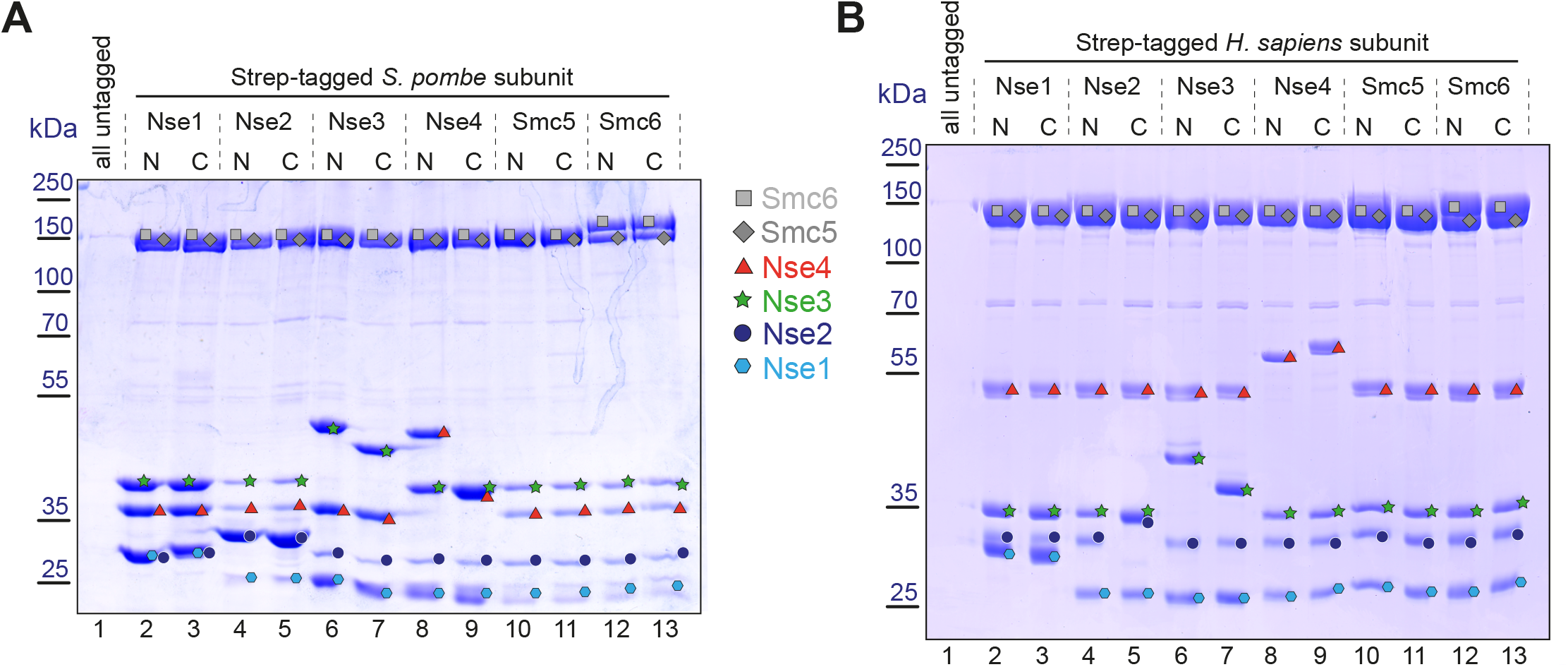
(A) Coomassie-stained SDS-gel showing the results of the expression screen for the *S. pombe* Smc5/6 complex in insect cells (Sf9) after infection with recombinant baculoviruses. For an overview of the cloning strategy for expression vectors see Fig. S3. Colored symbols for each subunit are used to identify the bands in the gel as in Fig. 2B.(B) Coomassie-stained SDS-gel showing the results of the expression screen for the *H. sapiens* Smc5/6 complex in mammalian cells. Subunits are again identified with colored symbols as in (A).

For expression of the human complex, we put the subunits under control of a CMV promoter and transfected the vectors resulting from the cloning procedure in **Fig. S3** into HEK293S cells (10 ml scale). Mammalian cell expression proved to be the most robust system, with the least variation regarding to placement of the affinity tag (**Fig. 4B**). Even though a slight excess of the Nse4/3/1 complex can be noticed in the gel when one of these subunits is tagged, this would likely not pose any problems because this excess of the smaller subunits/sub-complex can be removed by gel-filtration.

Our cloning strategy is thus easily adaptable to eukaryotic expression hosts and allows to quickly perform unbiased screens for ideal expression constructs in these systems.

## Discussion

We streamline the generation of expression construct variants for multi-subunit protein complexes with new vectors (**Table 1**) and protocols. 4G cloning eliminates time-consuming steps such as fragment amplification and purification, and using sequence-verified donors we create plasmids with up to four gene expression cassettes (GECs) in a single day in a sequence-conservative manner (**Fig. 1**).

Several points make 4G cloning unique and convenient: (i) ‘Donors’ for all GEC parts (promoters, tags, etc.) are sequence-verified only once, and can afterwards be freely combined without further sequencing of products. (ii) One single Type IIS restriction enzyme (BsaI) is required to carry out all described cloning steps, minimizing sequence domestication efforts. (iii) Handling of DNA fragments is kept to a minimum without time-consuming purification steps. (iv) Cloning of expression plasmids containing multiple GECs can be done in a single day with minimal hands-on-time. (v) Generation of many plasmid variants in parallel is straightforward and can be done with little additional effort. (vi) Using a simple protocol, any combination of ORFs can be fused with either ribosomal binding sites or peptide linkers for polycistronic expression or production of fusion proteins, respectively.

The fact that important regulatory regions and protein-coding sequences are protected from assembly-induced mutations saves additional time and makes the procedure more cost-efficient. For each new project only the new ORF-donors carrying individual subunits, and not the final products for expression screening need to be sequenced. Taking our case for the budding yeast Smc5/6 complex as an example, this required sequencing of only 10.5 kbp for the six ORF-donors shown in **Fig. 3A** and not the resulting >140 kbp of inserts from all 14 pMulti-plasmid versions. We have confirmed the sequence of selected expression constructs by whole-plasmid sequencing. Regulatory sequences and vector backbones are customizable, facilitating parallel construct creation for pro- and eukaryotic systems. Simultaneous transformation/transfection of several such multi-subunit vectors allows for the expression of even larger assemblies (>10 subunits). For eukaryotic expression we made our strategy compatible with biGBac vectors (Weismann), allowing generation of even larger vectors containing up to 20 subunits, but it should be noted that this option requires sequence-domestication of donors not only for BsaI but also for PmeI (3).

We validated our approach by screening various expression constructs for the hexameric Smc5/6 complex. Our results demonstrate that this eukaryotic assembly from yeasts and humans can be efficiently produced in *E. coli*, insect cells, or mammalian cells when all subunits are co-expressed. Our streamlined method enabled parallel generation of multiple plasmids in a single day for an unbiased screen to determine the optimal position of a specific affinity tag. While tag positioning was less critical for *S. pombe* and human complexes in insect or mammalian cells (Fig. 4), it was crucial for expressing the *S. cerevisiae* complex in *E. coli*, where only one option yielded a stoichiometric complex already after the affinity step. Once a suitable vector is cloned for wild-type complex expression, mutant versions of individual subunits can be created using the respective mutant donors. We extensively utilized this for the budding yeast complex to produce versions with ATPase and DNA-binding mutations or cysteine-substitutions for chemical cross-linking (11, 13). In addition to screening for the best position for a given tag, several different tags for a given subunit can also easily be compared, and we applied this to the JetABC complex from *E. coli*.

Establishing 4G cloning in a new laboratory requires some investments, mainly in getting accustomed to the procedure and getting hold of the donor and acceptor vectors, however, any efforts are quickly off-set by improved cloning and better material. Several projects in our laboratory have significantly benefited from 4G cloning.

## Acknowledgements

We are grateful to members of the Gruber lab for stimulating discussions and advice during the development of 4G cloning, to the Protein Production and Structure Core Facility at EPFL for protein expression, and to Kelvin Lau for helpful feedback on the manuscript.

## Materials and Methods

### Reagents

Gibson assembly mix is prepared as described in the original publication (9) with the exception that Taq DNA-Ligase is not included in the mixture. For generating circular plasmids for transformation, ligation of the nicks is not required, and removing the ligase significantly reduces the cost per reaction. For Golden Gate assembly BsaI HF-v2 enzyme (NEB) and T4-DNA Ligase (1u/µl; ThermoFisher) are used. Plasmids are maintained in the bacterial strains: DH5α (ThermoFisher) for standard plasmids, pirHC (Geneva Biotech) for donor plasmids containing the R6Kγ replication origin, and *ccdB*-Survival cells for empty pMulti plasmids containing the *ccdB* cassette. For expression of yeast proteins in *E. coli* we use the Rosetta strain, and for expression of the *E. coli* JetABC complex we employ BL21(DE3)Gold.

### Donor amplification for Gibson assembly

The donor vector (pD) is amplified and linearized for subsequent insertion of sequences by Gibson assembly by PCR using the primer pair pD_lin_fw (GGAGACCCACTGCTTGAGC) and pD_lin_rev (TGAGACCTAATATTCCGGAGTAG). 50 µl reactions contain 1x Phusion HF reaction buffer, 0.4 µM of each oligonucleotide, 0.4 mM dNTPs, 1 unit of Phusion DNA Polymerase (NEB), as well as 10 pg of a pD-template. After an initial denaturation step at 95°C for 1 min, 30 PCR cycles with 3 steps (95°C for 10 sec, 58°C for 10 sec, 72°C for 30 sec) produces the desired product. A 5 µl aliquot is mixed with DNA loading dye and analyzed on a 1 % agarose gel to verify amplification of the desired fragment. The product can be directly used for Gibson assembly, but it is recommended to purify it using a PCR purification kit (Qiagen) to be able to accurately measure the DNA concentration. After determining it, the DNA is diluted to a final concentration of 25 ng/µl.

### Creation of donor vectors for ORFs and tags

Coding sequences for protein targets as well as for N- and C-terminal tags are amplified by PCR with primers containing appropriate overhangs for subsequent Gibson assembly into the linearized donor plasmid. For creating ORF-donors these overhangs are 5-GGAATATTA**GGTCTC**A<u>CCAT</u>G-3’ for the forward primer and 5’-CAAGCAGTG**GGTCTC**C<u>ATCC</u>-3’. BsaI sites that flank the insert and are used for Golden Gate assembly are shown in bold, the four underlined nucleotides show the specific ‘sticky end’ created by BsaI digestion. In the forward primer the three nucleotides on the 3’ end (ATG) anneal to the start codon for the ORF. When creating an ORF for an N-terminal truncation of a protein, an ATG has to be added as the first codon. The reverse primer has to be designed in a way to exclude the native Stop-codon if C-terminal tagging should be an option. Coding sequences for N-terminal tags are amplified with 5’-CTCCGGAATATTA**GGTCTC**A<u>GGCG</u>-3’ overhangs on forward primers and 5’-GGCTCAAGCAGTG**GGTCTC**C<u>ATGG</u>C-3’ overhangs on reverse primers, while for C-terminal tags these overhangs are 5’-CTCCGGAATATTA**GGTCTC**A<u>GGAT</u>CC-3’ on forward and 5’-GGCTCAAGCAGTG**GGTCTC**C<u>CTTA</u>-3’ on reverse primers. If the sequence to be amplified does not contain internal BsaI sites, a single PCR using forward and reverse primers containing the overhangs described above is performed. In the case of internal sites, two additional primers and an additional PCR reaction are needed for each site (see **Fig. S1C**) to destroy these sites with silent mutations. PCR reactions contain the same components as described for linearization of the Donor vector, and specific cycling conditions (annealing temperature, elongation time) are chosen for each target. A 5 µl aliquot of the PCR product is analyzed on an agarose gel to verify that the desired fragment was obtained, that it is free of non-specific amplification products or primer-dimers, and to estimate product concentration. If only the desired product is visible, further purification is not necessary. While this makes it impossible to accurately determine the concentration using spectrophotometric methods, we find it sufficient to estimate the concentration based on agarose gel electrophoresis. If the desired fragment is not the only amplification product, the whole PCR reaction should be loaded on an agarose gel and the specific band purified using a gel extraction kit according to the manufacturer’s instructions.

### Insertion of sequences into the linearized donor by Gibson assembly

1 µl of linearized donor vector (at 25 ng/µl) is mixed with PCR product(s) of the insert corresponding roughly to a 2-5 x molar excess of insert in PCR tubes. Sterile water is added to 5 µl final volume and the tubes are put on ice. 5 µl of 2x Gibson-Mix are added on ice, and the tubes are quickly transferred to a pre-heated 50°C PCR block. After 15-30 min incubation the tubes are removed and placed on ice, and a 2 µl aliquot of the product is transformed into competent PIR1 *E. coli* cells (Invitrogen). The bacteria are plated on two LB-plates (for 90% or 10% of cells) containing 20 µg/ml chloramphenicol. After overnight incubation at 37°C colonies are picked and grown in 3 ml liquid cultures, and plasmids are isolated using a mini-prep kit (Qiagen). Cloned inserts are sequenced using the primers pD_seq_fw (5’-CGCGGTACCATAACTTCGTATAGC-3’) and/or pD_seq_rev (5’-GGGGTTATGATAGTTATTGCTCAGCGG-3’). Internal primers in the inserts are also used for long sequences.

### Linearization of multi-GEC acceptors for Gibson assembly

pMulti_ccdB plasmids are linearized by SwaI digestion in 40 µl reactions containing 50 ng/µl (roughly 15 nM) plasmid DNA, 1x NEB buffer 3.1, and 10 units of SwaI enzyme. After incubation at 25°C for 2 hours, 5 units of fresh enzyme are added followed by another 2 hours at 25°C. The enzyme is then inactivated by incubation at 75°C for 20 min. The material can be directly used as Gibson Acceptor without further purification because the *ccdB* cassette avoids cloning background due to incompletely digested plasmid. Linearization of pBig1 plasmids for making expression constructs for insect and mammalian cells was done by SwaI digestion following a previously published protocol (3).

### Generation of multi-GEC plasmids by 4G cloning

To create GECs from donor plasmids by Golden Gate assembly, 10 µl reactions in PCR tubes are assembled containing 1x T4 DNA Ligase buffer (ThermoFisher), 0.5 units of BsaIHF-v2, 0.25 units of T4 DNA Ligase, and 7.5 nM (1 µl each of a 75 nM stock) of all donors required to produce the desired GEC (donors for ORF, N- or C-terminal tag, promoter/terminator, as well as 5’ and 3’ Gibson overhang). Tubes are placed in a PCR machine and subjected to a cycling program (40 cycles of 37°C for 2 min and 16°C for 5 min, followed by a final step at 50°C for 10 min) to maximize the amount of fully assembled GECs. Compatible GECs created in this way are subsequently inserted by Gibson assembly into a suitable multi-GEC acceptor plasmid. For this, 2 µl of each GEC product are mixed with 1 µl of linearized multi-GEC acceptor (50 ng/µl) in PCR tubes on ice, the total volume is doubled using 2x Gibson-Mix, and tubes are quickly transferred to a PCR block pre-heated to 50°C. After incubation at this temperature for 30 min, 5 µl of the product are transformed into DH5α competent cells and bacteria are plated on LB-plates containing appropriate antibiotics.

### Expression tests in E.coli

Competent bacteria are transformed using expression vectors and plated on LB plates supplemented with the appropriate antibiotics. Colonies are inoculated in 10 ml terrific broth (TB) with the same antibiotics and grown to an OD(600nm) of about 1.0 at 37°C. The culture temperature is then decreased to 22°C and expression is induced by adding 0.5 mM of IPTG followed by overnight incubation at this lower temperature. Cells are harvested by centrifugation, resuspended in lysis buffer (50 mM Tris pH 7.5, 300 mM NaCl, 5 % glycerol), and sonicated with an MS73 probe (40 % output, 20 pulses of 1 s). The lysate is clarified by centrifugation at 16.000g at 4°C in a tabletop centrifuge, and added to StrepTactin Sepharose or Ni^2+^-NTA Sepharose (both from Cytiva) affinity resin that has been washed and equilibrated in lysis buffer. Tubes are incubated at 4°C on a rotating wheel for 1 hour, after which beads are pelleted by centrifugation (700g, 1 min) and washed twice with 1 ml lysis buffer. Bound material is eluted using 35 µl of lysis buffer supplemented with either 2.5 mM desthio-biotin (for StrepTactin) or 500 mM Imidazole (for Ni^2+^-NTA). After mixing the eluate with an equal volume of 2x SDS-loading dye, the sample is boiled for 5 min and proteins are separated on a 12 % SDS-PAGE gel followed by staining with Coomassie Brilliant Blue.

### Production of recombinant baculoviruses and protein expression tests in insect cells

Vectors for insect cell expression (pBig) are transformed into DH10 EMBacY cells and plated on LB plates containing Kanamycin, Gentamycin, Tetracyclin, IPTG, and X-Gal. White colonies are picked and grown in 4 ml of LB containing Kanamycin and Gentamycin. Cells are harvested and resuspended in 300 µl of buffer P1. 300 µl of buffer P2 are then added and the mixture is incubated at room temperature for 5 minutes. After addition of 300 µl of buffer N3 the mixture is incubated in ice for another 5 minutes before cell debris is removed by centrifugation at 4°C in an Eppendorf tabletop centrifuge. 800 µl of the supernatant are mixed with 600 µl of isopropanol and the tube is incubated at -20°C for 2 h to precipitate Bacmid DNA. After centrifugation for 10 minutes at 4°C at full speed in an Eppendorf tabletop centrifuge, the pellet is washed with 500 µl of 70 % ethanol and spun again. The ethanol is removed, and the pellet dried and resuspended in 50 µl of sterile water. 2 µl of this Bacmid DNA are transfected into Sf9 cells in a plate of a 6-well dish (0.8 x 10^6 cells per well), and the cells are incubated at 28°C. After 4 days the supernatants of these wells are harvested and 200 µl used to infect a 10 ml culture of Sf9 cells at a density of 10^6 cells/ml. After 3 days of incubation at 28°C (shaking at 180 rpm) the cells are harvested, and the pellets processed as described for small scale expression in *E*.*coli*.

### Expression tests in mammalian cell lines

Transfection of HEK293E cells with plasmid DNA (midi-prep scale) was carried out by the Protein Production and Structure Core Facility (PTPSP) at the Ecole Polytechnique Federal de Lausanne (EPFL). Briefly, HEK293E cells are grown in suspension in Ex-Cell medium (Sigma Aldrich) containing 4 mM Glutamine. For 10 ml of expression tests 10^7^ cells are harvested by centrifugation (1400 rpm for 6 min) and resuspended in RPMI 1640 medium (Invitrogen) containing 0.1 % pluronic at a concentration of 20 x 10^6^ cells /ml. 15 µg of plasmid DNA are added and mixed, followed by 30 µg of PEI-Max (Fisher Scientific). The mixture is then incubated for 90 minutes at 37°C with stirring, followed by dilution to 10^6^ cells/ml with Ex-Cell medium containing 4 mM Glutamine and transfer to Spin Tubes. Valproic acid (VPA) is added to a final concentration of 3.75 mM and the cultures are incubated for 3 days at 37°C. Cells are then harvested, the pellets washed once with PBS and frozen in dry ice. For lysis the pellets are thawed and resuspended in 2 ml of lysis buffer containing benzonase and cOmplete protease inhibitors (Roche), and processed as described for small scale expression in *E. coli*.

## References

1. Schütz, A., Bernhard, F., Berrow, N., Buyel, J.F., Ferreira-da-Silva, F., Haustraete, J., Heuvel, J. van den, Hoffmann, J.-E., Marco, A. de, Peleg, Y., et al. (2023) A concise guide to choosing suitable gene expression systems for recombinant protein production. STAR Protoc., 4, 102572.

2. Nie, Y., Chaillet, M., Becke, C., Haffke, M., Pelosse, M., Fitzgerald, D., Collinson, I., Schaffitzel, C. and Berger, I. (2016) Advanced Technologies for Protein Complex Production and Characterization. Adv. Exp. Med. Biol., 896, 27–42.

3. Weissmann, F., Petzold, G., VanderLinden, R., Veld, P.J.H. in ‘t, Brown, N.G., Lampert, F., Westermann, S., Stark, H., Schulman, B.A. and Peters, J.-M. (2016) biGBac enables rapid gene assembly for the expression of large multisubunit protein complexes. Proc. Natl. Acad. Sci., 113, E2564–E2569.

4. Qin, Y., Tan, C., Lin, J., Qin, Q., He, J., Wu, Q., Cai, Y., Chen, Z. and Dai, J. (2016) EcoExpress?Highly Efficient Construction and Expression of Multicomponent Protein Complexes in Escherichia coli. ACS Synth. Biol., 5, 1239–1246.

5. Fitzgerald, D.J., Berger, P., Schaffitzel, C., Yamada, K., Richmond, T.J. and Berger, I. (2006) Protein complex expression by using multigene baculoviral vectors. Nat. Methods, 3, 1021–1032.

6. Weber, E., Engler, C., Gruetzner, R., Werner, S. and Marillonnet, S. (2011) A Modular Cloning System for Standardized Assembly of Multigene Constructs. PLoS ONE, 6, e16765.

7. Engler, C., Gruetzner, R., Kandzia, R. and Marillonnet, S. (2009) Golden Gate Shuffling: A One-Pot DNA Shuffling Method Based on Type IIs Restriction Enzymes. PLoS ONE, 4, e5553.

8. Engler, C., Kandzia, R. and Marillonnet, S. (2008) A One Pot, One Step, Precision Cloning Method with High Throughput Capability. PLoS ONE, 3, e3647.

9. Gibson, D.G., Young, L., Chuang, R.-Y., Venter, J.C., Hutchison, C.A. and Smith, H.O. (2009) Enzymatic assembly of DNA molecules up to several hundred kilobases. Nat. Methods, 6, 343–345.

10. Halleran, A.D., Swaminathan, A. and Murray, R.M. (2018) Single Day Construction of Multigene Circuits with 3G Assembly. ACS Synth. Biol., 7, 1477–1480.

11. Taschner, M. and Gruber, S. (2023) DNA segment capture by Smc5/6 holocomplexes. Nat. Struct. Mol. Biol., 30, 619–628.

12. Liu, H.W., Roisné-Hamelin, F., Beckert, B., Li, Y., Myasnikov, A. and Gruber, S. (2022) DNA-measuring Wadjet SMC ATPases restrict smaller circular plasmids by DNA cleavage. Mol. Cell, 82, 4727-4740.e6.

13. Taschner, M., Basquin, J., Steigenberger, B., Schäfer, I.B., Soh, Y., Basquin, C., Lorentzen, E., Räschle, M., Scheltema, R.A. and Gruber, S. (2021) Nse5/6 inhibits the Smc5/6 ATPase and modulates DNA substrate binding. EMBO J., 40, e107807.

14. Roisné-Hamelin, F., Liu, H.W., Taschner, M., Li, Y. and Gruber, S. (2024) Structural basis for plasmid restriction by SMC JET nuclease. Mol. Cell, 84, 883-896.e7.

15. Haering, C.H. and Gruber, S. (2016) SnapShot: SMC Protein Complexes Part I. Cell, 164, 326-326.e1.

16. Yatskevich, S., Rhodes, J. and Nasmyth, K. (2019) Organization of Chromosomal DNA by SMC Complexes. Annu. Rev. Genet., 53, 1–38.

17. Aragón, L. (2018) The Smc5/6 Complex: New and Old Functions of the Enigmatic Long-Distance Relative. Annu. Rev. Genet., 52, 89–107.

18. Bermúdez-López, M., Pociño-Merino, I., Sánchez, H., Bueno, A., Guasch, C., Almedawar, S., Bru-Virgili, S., Garí, E., Wyman, C., Reverter, D., et al. (2015) ATPase-Dependent Control of the Mms21 SUMO Ligase during DNA Repair. PLoS Biol., 13, e1002089.

19. Zabrady, K., Adamus, M., Vondrova, L., Liao, C., Skoupilova, H., Novakova, M., Jurcisinova, L., Alt, A., Oliver, A.W., Lehmann, A.R., et al. (2016) Chromatin association of the SMC5/6 complex is dependent on binding of its NSE3 subunit to DNA. Nucleic Acids Res., 44, 1064–1079.

20. Deep, A., Gu, Y., Gao, Y.-Q., Ego, K.M., Herzik, M.A., Zhou, H. and Corbett, K.D. (2022) The SMC-family Wadjet complex protects bacteria from plasmid transformation by recognition and cleavage of closed-circular DNA. Mol. Cell, 82, 4145-4159.e7.

21. Liu, H.W., Roisné-Hamelin, F. and Gruber, S. (2023) SMC-based immunity against extrachromosomal DNA elements. Biochem. Soc. Trans., 51, 1571–1583.

22. Pradhan, B., Deep, A., König, J., Baaske, M.D., Corbett, K.D. and Kim, E. (2024) Loop extrusion-mediated plasmid DNA cleavage by the bacterial SMC Wadjet complex. bioRxiv, 10.1101/2024.02.17.580791.

