## Supplementary Figures for "4G cloning: rapid gene assembly for expression of multisubunit protein complexes in diverse hosts"

**Supplemental Figure legends**

Figure S1:

(A) Schematic representation of different types of element donors which differ with respect to the overhangs created at their 5' and 3' ends.

(B) Creation of a donor plasmid by PCR and Gibson assembly into pD. The target sequence is amplified with primers containing BsaI restriction sites and the desired 'sticky ends' for the respective donor type. The resulting PCR product is then inserted into PCR-linearised pD by Gibson assembly.

(C) Creation of a donor plasmid by PCR and Gibson assembly into pD, if the target contains an internal BsaI site. As in (B) but two additional primers are required to introduce a silent mutation in the internal BsaI-site.

(D) BsaI 'sticky' ends on the 5' and 3'-ends of donors for ORF-elements as well as N- and C-terminal TAG-elements in their sequence contexts.

(E) BsaI 'sticky' ends created by promoter/terminator ('P+T') donors around promoters (black triangles) and terminators (black rounded rectangles). Each donor exists in 3 versions to accommodate untagged ORFs, or N- or C-terminally tagged ORFs.

(F) BsaI 'sticky' ends created by Gibson donors. Each Gibson donor contains a homology sequence used for assembly, which is separated from the rest of the GEC by 200-300 bp spacers.

(G) Overview on the cloning of the efficiency test shown in Fig 2. Individual GECs were assembled fro 4 ORFs (ScNse4, ScNse3, ScNse2, ScNse1), and T7 promoters and terminators were chosen as flanking regulatory elements. For all but ScNse1 two versions were created differing in their terminal Gibson overhangs. Compatible fragments were combined in separate tubes with linearized acceptor (pMulti_A) and expression plasmids were created by Gibson assembly.

Figure S2:

Schematic overview of the procedure described in Fig. 3A with all intermediate fragments and vectors. Each ORF donor is used to create three versions of either untagged, N- or C-terminally tagged GECs with specific Gibson overhangs, and those are used to create 7 pMulti_K plasmids (containing Nse4, Nse3, and Nse1) or 7 pMulti_A plasmids (containing Smc6, Smc5, and Nse2). These 14 plasmids are then transformed in 13 combinations (all untagged or 12 combinations with each individual affinity tag position) into *E. coli* and expression tests are carried out with StrepTactin pulldowns.

Figure S3:

Schematic overview of the procedure described in Fig. 4 with all intermediate fragments and vectors. Each ORF donor was used to created three versions of either untagged, N- or C-terminally tagged GECs with specific Gibson overhangs. *S. pombe* subunits were given polyhedrin promoters, and *H. sapiens* subunits obtained CMV promoters. Those fragments were used to create 7 pBig1a plasmids (containing Nse4, Nse3, and Nse1) or 7 pBig1b plasmids (containing Smc6, Smc5, and Nse2). Plasmids containing all subunits were created from pBig1a and pBig1b following the procedure described by Weissmann *et al*. Expression tests were carried out in insect or mammalian cells following standard procedures.

### SUPPLEMENTAL FIGURE 1

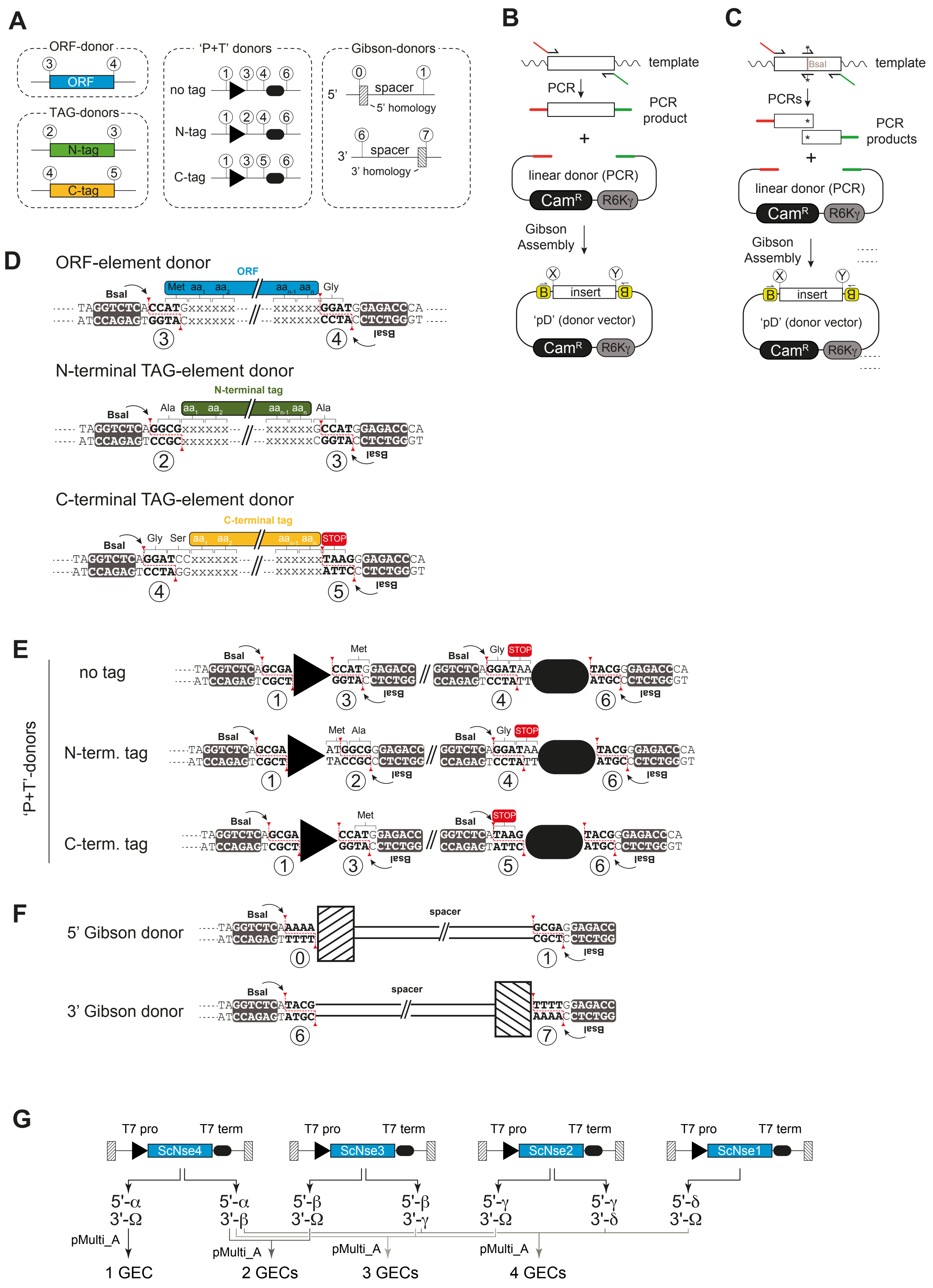

### SUPPLEMENTAL FIGURE 2

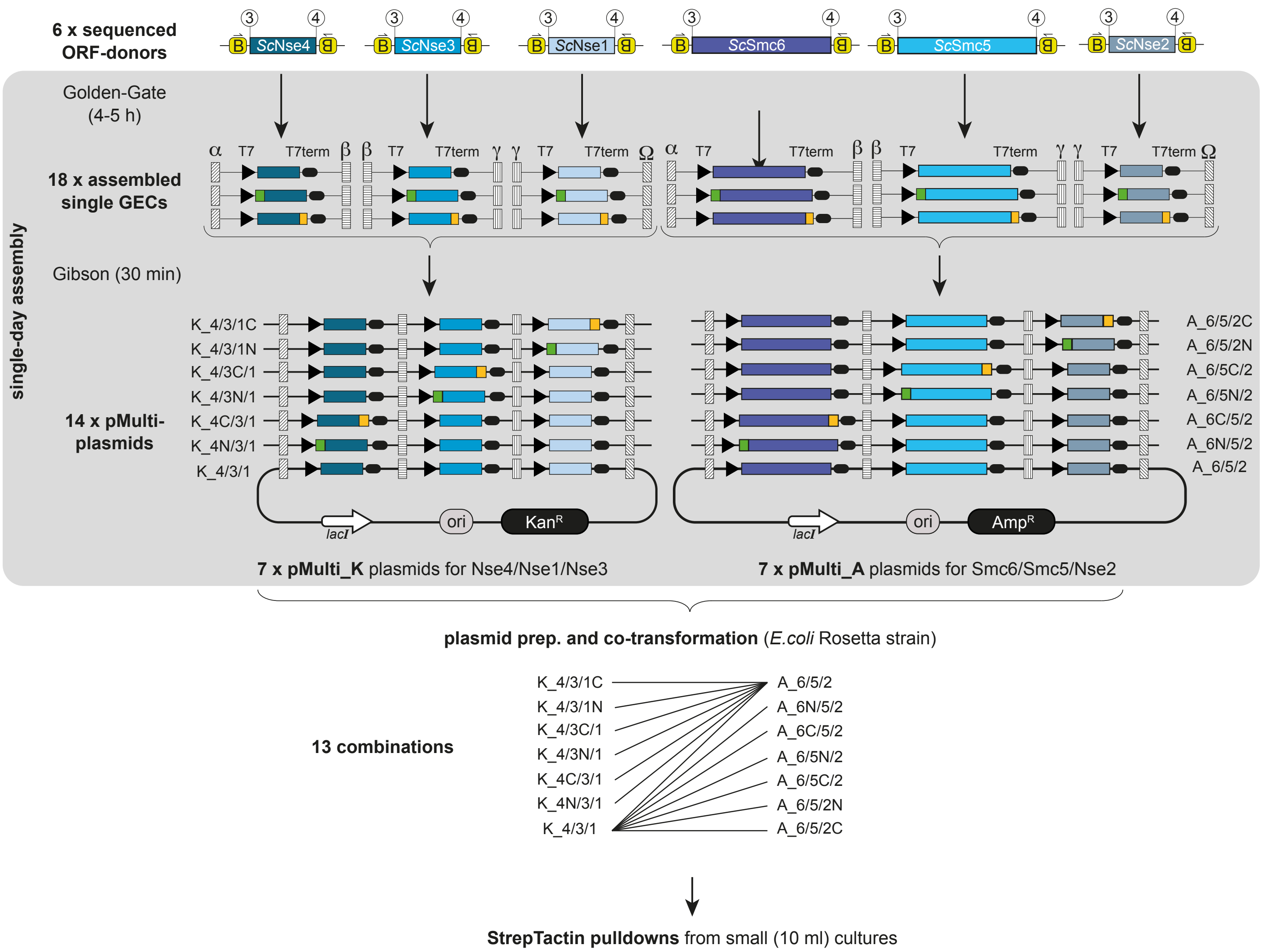

### SUPPLEMENTAL FIGURE 3

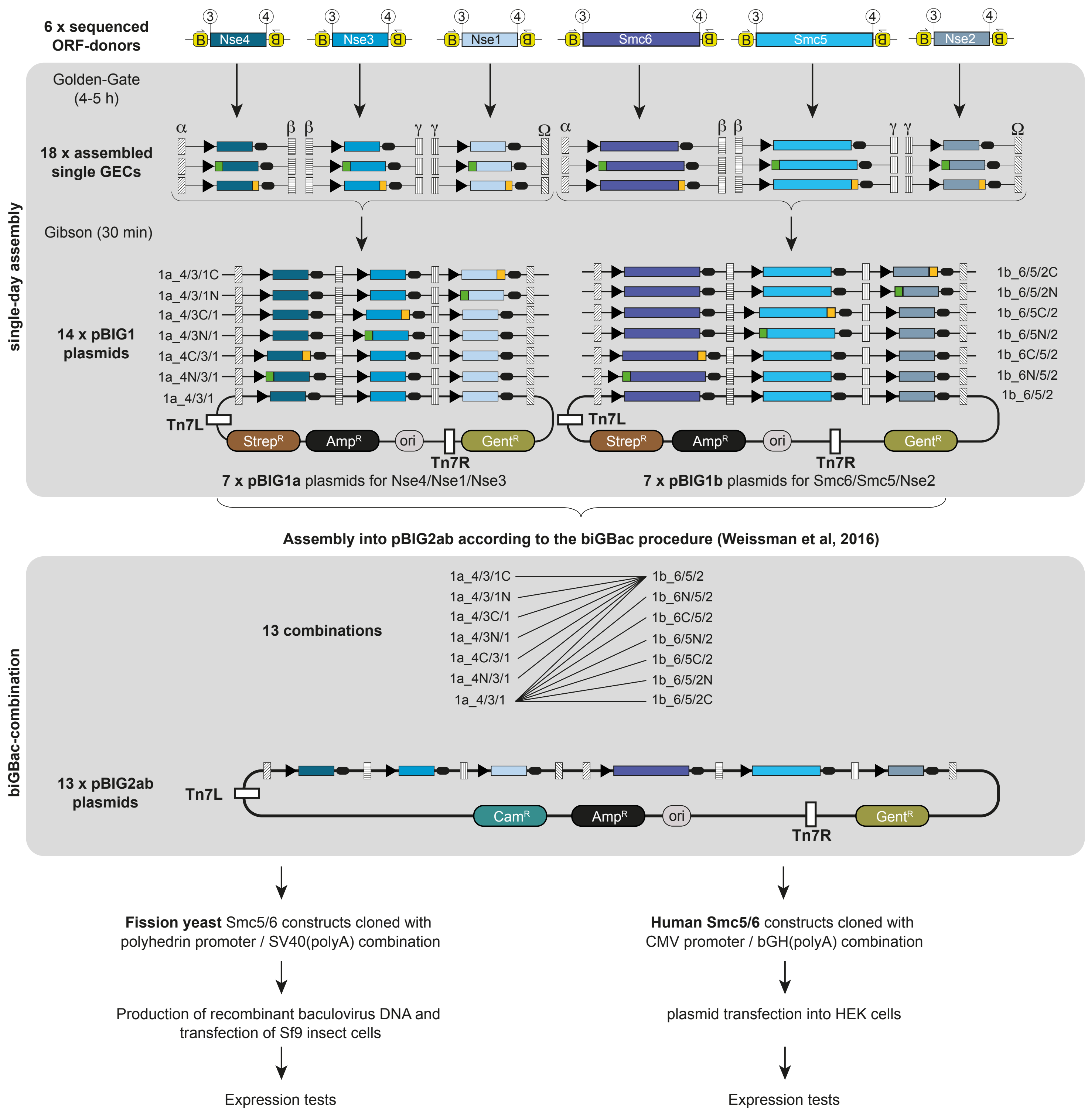
